# Alzheimer’s Disease Patient Brain Extracts Induce Multiple Pathologies in Vascularized Neuroimmune Organoids for Disease Modeling and Drug Discovery

**DOI:** 10.1101/2024.10.28.620680

**Authors:** Yanru Ji, Xiaoling Chen, Meek Connor Joseph, Jenna Lillie McLean, Yang Yang, Chongli Yuan, Jean-Christophe Rochet, Fei Liu, Ranjie Xu

## Abstract

Alzheimer’s Disease (AD) is the most common cause of dementia afflicting 55 million individuals worldwide, with limited treatment available. Current AD models mainly focus on familial AD (fAD), which is due to genetic mutations. However, models for studying sporadic AD (sAD), which represents over 95% of AD cases without specific genetic mutations, are severely limited. Moreover, the fundamental species differences between humans and animals might significantly contribute to clinical failures for AD therapeutics that have shown success in animal models, highlighting the urgency to develop more translational human models for studying AD, particularly sAD. In this study, we developed a complex human pluripotent stem cell (hPSC)-based vascularized neuroimmune organoid model, which contains multiple cell types affected in human AD brains, including human neurons, microglia, astrocytes, and blood vessels. Importantly, we demonstrated that brain extracts from individuals with sAD can effectively induce multiple AD pathologies in organoids four weeks post-exposure, including amyloid beta (Aβ) plaques-like aggregates, tau tangles-like aggregates, neuroinflammation, elevated microglial synaptic pruning, synapse/neuronal loss, and impaired neural network. Furthermore, after treatment with Lecanemab, an FDA-approved drug targeting Aβ, AD brain extract exposed organoids showed a significant reduction of amyloid burden. Thus, the neuroimmune organoid model provides a unique opportunity to study AD, particularly sAD under a pathophysiological relevant three-dimensional (3D) human cell environment. It also holds great promise to facilitate AD drug development, particularly for immunotherapies.

## Introduction

Alzheimer’s Disease (AD) is a devastating neurodegenerative disorder and the most common cause of dementia, affecting more than 55 million individuals globally[1, 2]. Despite extensive efforts, most therapeutics have failed in clinical trials till now, with only limited effective treatments currently available[3–5]. AD is manifested with progressive cognitive decline and is characterized by multiple pathological hallmarks, including extracellular amyloid beta (Aβ) plaques, intracellular microtube-associated tau neurofibrillary tangles (NFTs), neuroinflammation, synapse/neuronal loss, and brain atrophy[2, 4, 6]. To understand the mechanisms and develop therapeutics for AD, various animal models have been developed and significantly advanced our understanding of the disease mechanism[7–10]. However, most existing models focus on dominantly inherited familial AD (fAD), which is caused by genetic mutations in genes identified in individuals with fAD, including amyloid precursor protein (APP), presenilin1 (PSEN1), and presenilin2 (PSEN2)[7, 10, 11]. In contrast, models for sporadic AD (sAD), which do not involve highly penetrating genetic mutations and account for 95% of all AD cases, are severely limited[3, 10]. While both fAD and sAD share similar pathological hallmarks, such as Aβ plaques and NFTs, the etiology of sAD is still largely unknown and may involve more complicated genetic and environmental interactions[9, 10]. Thus, developing appropriate models for sAD without genetic mutations is critically needed. Moreover, significant species difference exists between humans and rodents. For instance, many repeat sequences in the genome and gene splicing patterns are unique to humans[12]. These species differences may have profound impacts when modeling neurological disorders, particularly for AD, and may significantly contribute to the failures of many AD therapeutics that succeeded in pre-clinical animal studies but failed in clinical trials[3, 8, 13]. Therefore, it is critical to use human samples and develop human AD models that could recapitulate sAD features to advance our understanding of the disease mechanism and facilitate the development of therapeutic interventions.

Given the challenge of accessing and manipulating functional human brain tissues, human pluripotent stem cells (hPSCs), which include human embryonic stem cells (hESCs) and induced pluripotent stem cells (iPSCs), have emerged as valuable tools to utilize human cells for studying human neurological disorders, including AD[3, 14, 15]. The hPSC-based *in vitro* models, including two-dimensional (2D) and 3D models with pure human neural cells, are easy to manipulate and enable a high throughput for screening making them valuable for modeling AD[3, 16–18]. The hPSC 2D model is useful for dissecting the impact of genetic and other factors on the individual cell types involved in AD at a basic functional level[19–27]. On the other hand, recent studies have also highlighted the value of hPSC 3D models, such as brain organoids and multiple cell type co-culture-based models, which incorporate various neural cell types and intricate cell-cell and cell-matrix interactions and preliminary neural structures, offering a more physiologically relevant environment to model diverse aspects of AD[28–36]. For example, fAD iPSC-derived brain organoids have successfully replicated AD pathological events, including Aβ aggregates and hyperphosphorylated tau, and facilitated the understanding of AD, such as the roles of APOE4 and C3 complement in AD[28, 30, 33, 34].

However, critical limitations persist, impeding these 3D models from effectively modeling AD. (1) Most research used simple organoids composed mainly of neurons and astrocytes. Although some studies have integrated microglia into organoids, or incorporated blood vessels into 3D neural culture, it is still a challenge to integrate multiple cell types, particularly microglia and blood vessels simultaneously within one organoid to reflect the complex pathophysiological environment of AD brains. (2) The current organoid models for AD study mainly focused on fAD, and these models usually take three to six months to exhibit AD pathologies[30, 34, 35, 37]. An intriguing recent study induced sAD-like phenotypes by exposing healthy organoids to serum to mimic the condition of blood-brain barrier (BBB) leakage, a risk factor for sAD[36]. However, a brain organoid model that contains multiple cell types affected in AD brains, capable of effectively recapitulating key AD pathological hallmarks of sAD within a relatively short time frame induced by an efficient trigger for modeling sAD, is still lacking.

To circumvent these limitations, in this study, we developed a vascularized neuroimmune organoid model for studying AD, particularly for sAD. This organoid model contains multiple cell types that are affected in human AD brains, including human neurons, astrocytes, microglia, and blood vessels. Since AD postmortem brain tissues contain proteopathic seeds, including Aβ and tau, that have the prion-like seeding activity to induce counterpart normal protein aggregate in animal models[38–42], we hypothesize that sAD postmortem tissues-derived brain extracts that contain both Aβ and tau seeds, can induce multiple AD pathologies in human cells in organoids, mirroring observations in sAD. Remarkably, our results demonstrated that organoids exposed to AD brain extracts successfully recapitulated multiple AD-like pathologies, including Aβ plaque-like aggregates, tau tangle-like aggregates, neuroinflammation, elevated microglial synaptic pruning, synapse/neuronal loss, but not organoids treated with vehicle control. Furthermore, we validated our organoids for drug discovery using Lecanemab, an FDA-approved anti-Aβ antibody for AD treatment, and observed a significant alleviation of Aβ burden after treatment in the organoids that have been exposed to AD brain extracts. Thus, this vascularized neuroimmune organoid model presents a unique opportunity to study sAD and holds great promise to facilitate AD drug development.

## Results

### Generation and characterization of vascularized neuroimmune organoids

To develop a brain organoid model that contains major cell types affected in human AD brains, building upon our previously established microglia-containing brain organoid model[43], we aim to introduce blood vessels into these organoids and develop a vascularized neuroimmune organoid model by co-culture of hPSCs-derived neural progenitor cells (NPCs), primitive macrophage progenitors (PMPs), and vascular progenitors (VPs) under 3D condition. To achieve this goal, we first generated NPCs and PMPs following our previous studies[43–45] (**Figure 1A**). The identity of hPSCs-derived NPCs was confirmed by co-expression of neural progenitor cell markers, PAX6 and NESTIN. PMPs were confirmed by co-expression of CD235, a marker for YS primitive hematopoietic progenitors[43], and CD43, a marker for hematopoietic progenitor-like cells[43] (**Figure 1B**). VPs were generated using a published protocol with slight modifications detailed in Materials and Methods[46]. To generate brain organoids, the initial number was calibrated and co-cultured at 30,000 for NPCs, 12,000 for PMPs, and 7,000 for VPs, based on our previous experience. These cells spontaneously assembled and formed organoids on day 1. Subsequent culture of the organoids was supplemented with mitogen fibroblast growth factor (bFGF) for 5 days to promote cellular proliferation (referred to as proliferation stage). Thereafter, organoids were transitioned to a neural differentiation medium which contained neurotrophic factors, interleukin-34 (IL-34), vascular endothelial growth factor (VEGF), and other necessary supplements for long-term culture to support neuronal, microglial, and vascular maturation (referred to as Differentiation stage) (**Figure 1A**). To ensure genetic diversity and robustness of our findings, three different hPSC lines (two iPSC lines, and one hESC line) from healthy individuals were used to derive progenitor cells.

**Figure 1.**
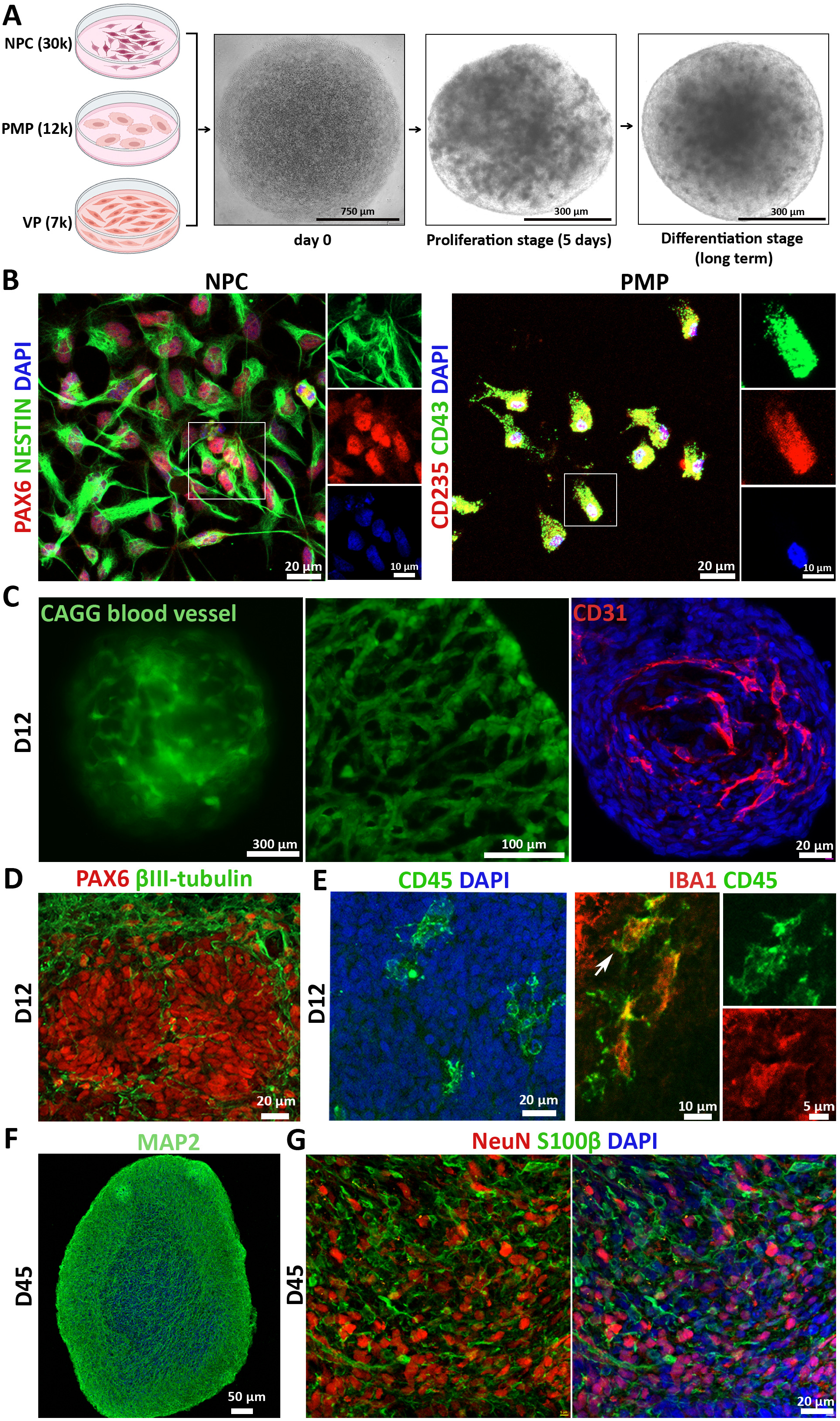
Generation and characterization of vascularized neuroimmune organoids. (A) Schematic image of the generation of vascularized neuroimmune organoids. hPSCs-derived NPCs, PMPs, and VPs were co-cultured to form 3D spheroids. The proliferation stage lasted for five days before organoids were cultured in a differentiation medium. Scale bars, 750 or 300 µm as indicated. (B) Left panel, representatives of PAX6^+^ and NESTIN^+^ NPCs. Right panel, CD235^+^ and CD43^+^ PMPs. Scale bars, 20 or 10 μm as indicated. (C) Representatives of blood vessels in 12-day-old organoids. Left panel, CAGG-derived blood vessels in living organoids; Middle panel, sections of organoids incorporated with CAGG-derived blood vessels; Right panel, representative of CD31 staining. Scale bars, 300, 100, or 20 μm as indicated. (D) Representative of neural rosette stained with PAX6 and βIII-tubulin in 12-day-old organoids. Scale bar, 20 μm. (E) Characterization of microglia in 12-day-old organoids. Left panel, representative of CD45^+^ microglia; Right panel, representative of CD45^+^Iba1^+^ microglia. Scale bars, 20, 10, or 5 μm as indicated. (F) Representative of MAP2 expression in 45-day-old organoids. Scale bar, 50 μm. (G) Representatives of NeuN^+^ neurons and S100β^+^ astrocytes in 45-day-old organoids. Scale bar, 20 μm.

To visualize the formation of blood vessels in living organoids, we incorporated VPs from an additional GFP-expressing hESC line (CAGG line). This allows us to track the vascular lineage cells by GFP signal. Under live cell imaging, clear and distinct branching structures were revealed in organoids on day 12 of culture (**Figure 1C**, left panel). Organoid sections without immunostaining also displayed lumen-like structures (**Figure 1C**, middle panel, and **Figure S1**). We further stained organoids with CD31, an endothelial cell marker, and observed CD31-expressing cells in organoids. The results confirmed the establishment of vascular structures within organoids (**Figure 1C**, right panel). At day 12, ventricular zone-like regions were seen in organoids, which contained PAX6+ progenitors and βIII-Tubulin+ immature neurons (**Figure 1D**), mimicking the proliferative region in the developing cortical brain. To confirm PMPs differentiate into microglia at the differentiation stage, we first examined the expression of CD45 at day 12, which is a marker for all nucleated hematopoietic cells[43]. CD45+ cells exhibited a relatively even spatial distribution in organoids (**Figure 1E**, left panel). Microglial identity was further confirmed by double staining of CD45 and IBA1, a canonical macrophage/microglia marker[43]. CD45 and IBA1 double staining showed ramified morphology, indicating healthy and functional microglia residing in organoids (**Figure 1E**, right panel). Previous studies have shown that NPCs can mature into neuronal lineage cells, including neurons and astrocytes in brain organoids[43, 47]. To confirm neural maturation in the organoids, we first stained MAP2, a marker for neuronal dendrite, and observed robust expression of MAP2 at Day 45 (**Figure 1F**), suggesting NPCs efficiently differentiated into neurons. Further staining with NeuN, a mature neuron marker, and S100β, an astrocyte marker, revealed robust NeuN+ cells and S100β+ cells, demonstrating that NPCs differentiated into mature neurons and astrocytes during the differentiation stage (**Figure 1G**). Taken together, we successfully developed vascularized neuroimmune organoids incorporating mature neurons, microglia, and vascular networks, which allows us to model the complex human brain environment.

### Sporadic AD (sAD) patient-derived brain extracts induce amyloid pathology in organoids

Our next aim is to enable the vascularized neuroimmune organoids to efficiently recapitulate key AD pathological features for studying sAD without genetic mutations. Prior studies have shown that human misfolded proteopathic seeds, such as Aβ or tau, can induce corresponding pathologies in transgenic mice engineered to express these human proteins[38–42]. Notably, recent studies suggested that Aβ seeds may also be able to transmit among humans, for instance, individuals who received Aβ contaminated cadaver-derived pituitary growth hormone (c-hGH) during childhood developed Aβ pathologies and cerebral amyloid angiopathy (CAA) later in their life[48–50]. Based on these findings, we hypothesized that AD individual-derived brain extracts that contain proteopathic seeds, including Aβ or tau, will induce AD pathologies in organoids as seen in human AD brains. To test our hypothesis and model sAD, we treated organoids with sAD individual postmortem tissue-derived brain extracts (henceforth, the AD group) to model sAD. The corresponding vehicle (buffer) was used as control (henceforth, the vehicle group). In addition, age-matched healthy individual-derived brain extracts were also included. Organoids after 10 days of neuronal differentiation were exposed to AD brain extracts or the vehicle for two days, and samples were collected at 2 weeks or 4 weeks post-exposure. Subsequently, we examined AD pathological hallmark expression, including Aβ and tau pathologies, inflammation, and synapse/neuronal loss in organoids (**Figure 2A**).

**Figure 2.**
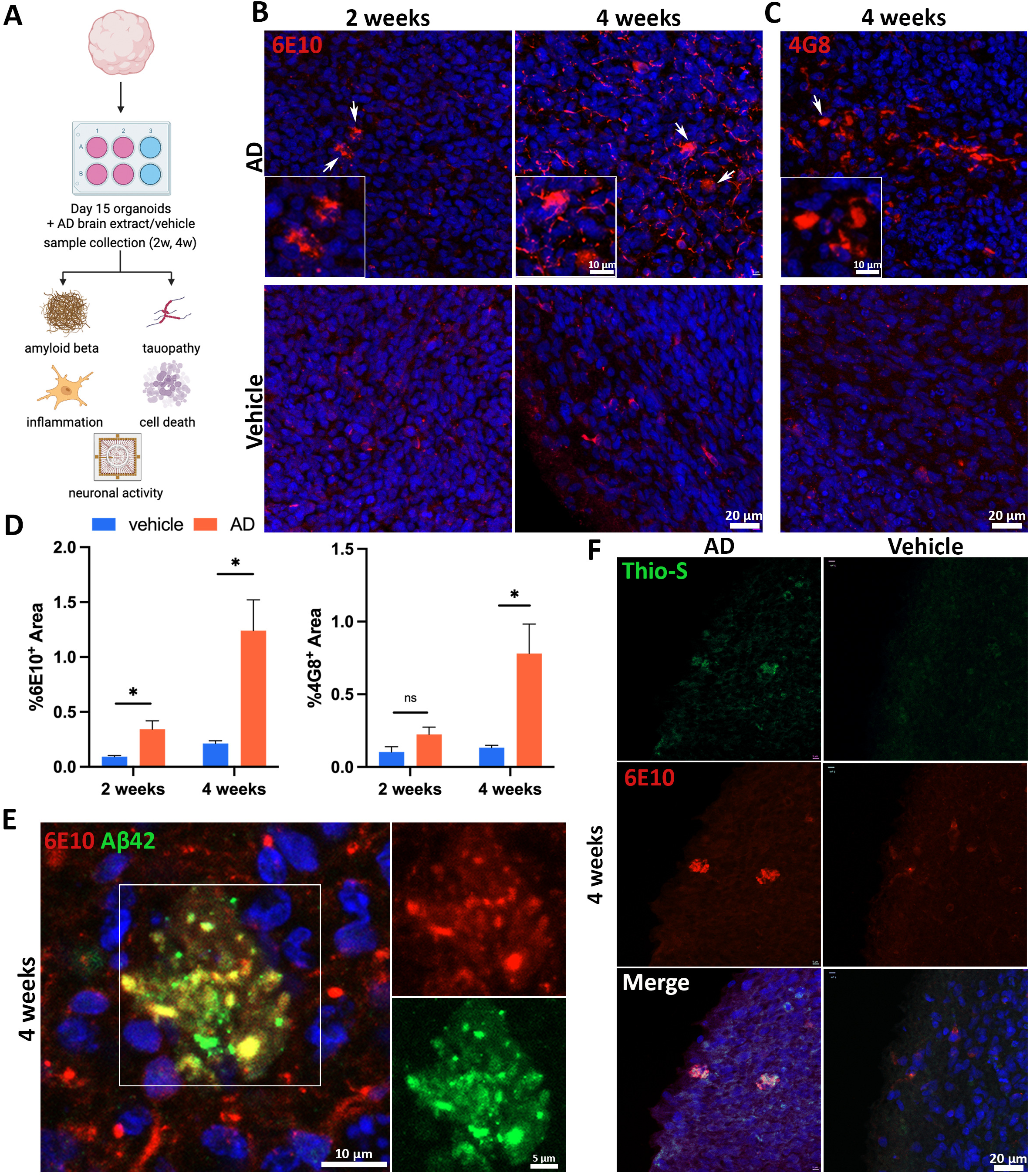
Sporadic AD patient-derived brain extracts induce amyloid pathology in organoids. (A) Schematic image of experimental design. (B) Representatives of 6E10 in organoids at 2 or 4 weeks post-exposure to AD brain extracts or vehicle. Scale bars, 20 or 10 µm as indicated. (C) Representatives of 4G8 in organoids at 2 or 4 weeks post-exposure to AD brain extracts or vehicle. Scale bars, 20 or 10 µm as indicated. (D) Quantification of percentage area of 6E10^+^ signals and 4G8^+^ signals over time. n=4 independent experiments from 3 hPSC lines, each experiment used one hPSC line and contained 4-6 organoids. Data are presented as mean ± SEM. Unpaired t test with Welch’s correction, *p < 0.05, ns represents no significance. (E) Representatives of co-staining with 6E10 and Aβ42 in organoids at 4 weeks post-exposure to AD brain extracts. Scale bars, 10 or 5 µm as indicated. (F) Representatives of Thioflavin-S^+^ and 6E10^+^ structures in organoids at 4 weeks post-exposure to AD brain extracts or vehicle. Scale bar, 20 μm.

To examine Aβ pathology characterized by extracellular Aβ aggregates in organoids, we utilized 6E10 and 4G8 antibodies, which can detect the amino acid residues 3-8 or 17-24 of Aβ, respectively[51]. Two weeks post brain extract or vehicle exposure, a modest number of weakly diffuse 6E10 and 4G8 signals were detected in both the AD and vehicle groups (**Figures 2B and S2**). Interestingly, only the AD group exhibited intraneuronal Aβ aggregates, as indicated by the presence of discrete, punctate 6E10 signals around nuclei (**Figure 2B**, top panel). To exclude the possibility that aggregates were caused by other components contained in the brain extracts other than toxic protein seeds, age-matched healthy individual-derieved brain extracts were included as the healthy control group. Similar to the vehicle group, diffuse but not aggregated signals were observed in healthy control brain extract-treated organoids (**Figure S3**). Since 6E10 can detect both Aβ and intraneuronal amyloid precursor protein (APP)[51], the diffuse 6E10 and 4G8 signals might represent the endogenous APP. To confirm the presence of intraneuronal Aβ aggregates in the AD group, we stained organoids with MOAB-2 antibody, which can specifically label intraneuronal Aβ, but not APP[51]. MOAB-2-expressing cells were only detected in the AD group, confirming the formation of intraneuronal Aβ aggregates (**Figure S4**).

Remarkably, after 4 weeks post AD brain extract exposure, multiple dense extracellular plaque-like aggregates were detected in the AD group but not the vehicle group(**Figure 2B**) or healthy control brain extract treated organoids (**Figure S3**). These extracellular aggregates were consistently identified in the AD group by 4G8 staining (**Figure 2C**). Quantitative analysis showed that compared to the vehicle group, the AD group demonstrates a significant escalation of Aβ burden, as indicated by the positive areas of 6E10 (AD vs Vehicle, 1.24 ± 0.28 vs 0.21 ± 0.03) and 4G8 (AD vs Vehicle, 0.78 ± 0.20 vs 0.13 ± 0.02) (**Figure 2D**). To confirm the composition of extracellular Aβ aggregates, we performed double staining of organoids with 6E10 with another antibody Aβ42, which primarily targets Aβ42[52]. The overlapped punctate signals of 6E10 and Aβ42 were observed from extracellular regions in the AD group, suggesting that extracellular Aβ aggregates contain Aβ42 (**Figure 2E**). Moreover, to assess Aβ plaque formation, we conducted double staining of organoids with 6E10 with a dye, Thioflavin-S, which binds to β-sheet structure in protein aggregates, including plaques and tangles[53]. Consistently, the vehicle group showed some diffuse intraneuronal 6E10 signals, which were Thioflavin-S negative; only the AD group exhibited the co-localization of strong extracellular 6E10 and Thioflavin-S signals, reinforcing the presence of plaque-like aggregates in AD group (**Figure 2F**, top panel). Collectively, these findings demonstrated that AD brain extracts containing proteopathic Aβ seeds can induce Aβ pathology, including Aβ plaque-like aggregates, in healthy brain organoids within a 4-week post-exposure time frame.

### Sporadic AD (sAD)patient-derived Brain extracts induce tau pathology in organoids

Tau pathology, caused by the abnormal accumulation of the microtubule-associated protein tau, is a major pathological hallmark of AD. In human AD brains, tau undergoes abnormal hyperphosphorylation and other modifications, which convert it into a pathological protein with prion-like seeding activity and form neurofibrillary tangles (NFTs)[54, 55]. Given that the AD brain extracts applied in this study contained tau seeds (**Figure S5**), we wonder whether the AD brain extract can induce tau pathology in our organoid model. To examine tau pathology, we utilized a widely used antibody, AT8 that can detect phosphorylated tau[41]. Notably, 2 weeks post-exposure, the AD group presented with puncta-like, AT8-positive intraneuronal signals (**Figure 3A**, top panel), potentially indicating the formation of aggregates in neurons. Notably, 4 weeks post-AD brain extract exposure, the intensity and quantity of condensed AT8 signals increased significantly, suggesting further formation of tau aggregates (**Figure 3A**, top panel). In contrast, the vehicle group displayed no such phosphorylated tau aggregates at either time point (**Figure 3A**, bottom panel), nor did healthy control brain extract treated organoids (**Figure S6**). Additionally, different from the condensed AT8 signals after AD brain extract exposure, weak and diffuse axonal AT8 positive signals were observed in organoids at 2 weeks post-exposure to the vehicle and healthy control brain extracts (**Figure 3A**, **S6, and S7**), aligning with prior studies that identified transient phosphorylated tau in neurons during development [48, 56].

**Figure 3.**
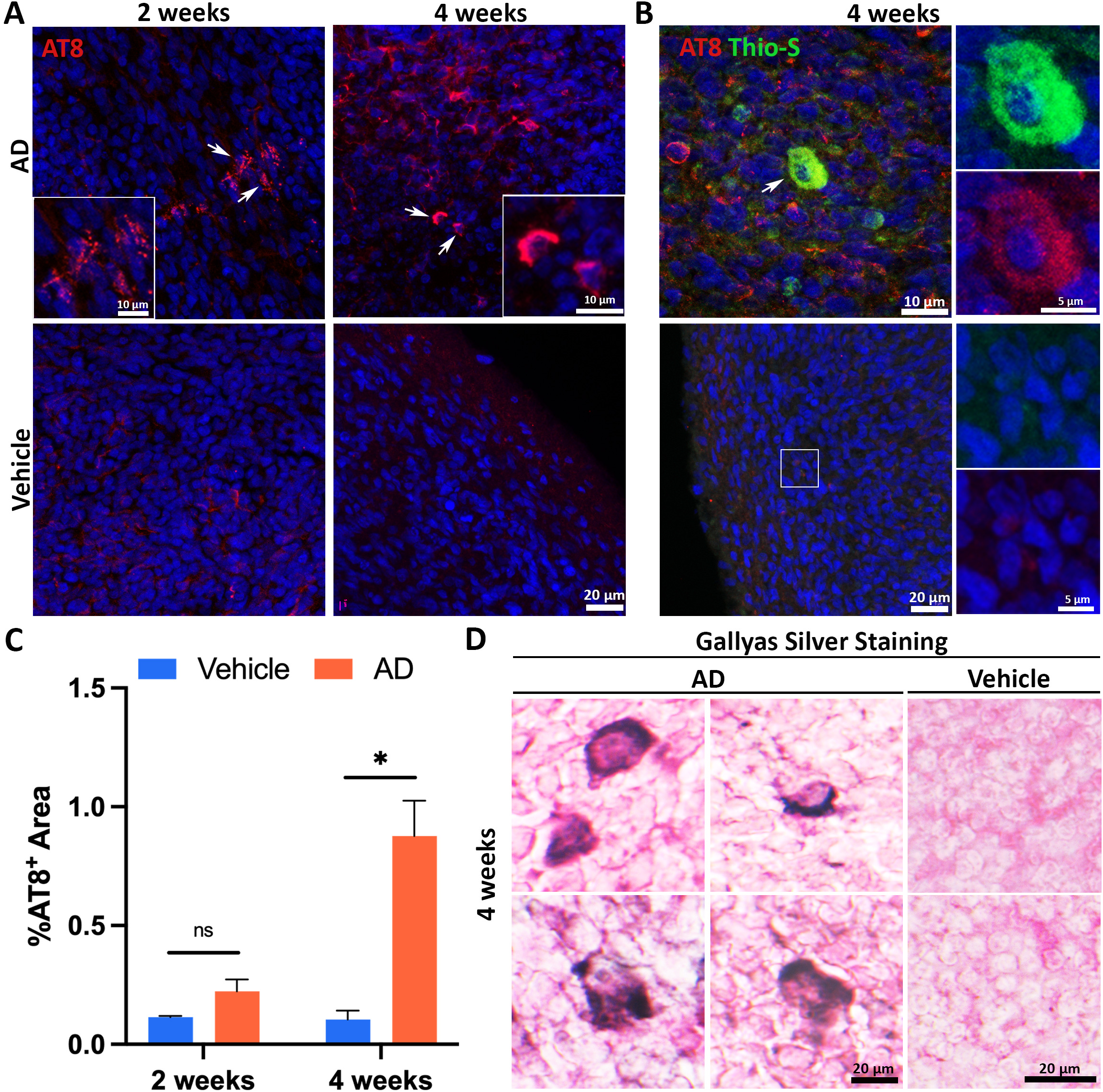
Sporadic AD patient-derived brain extracts induce tau pathology in organoids. (A) Representatives of AT8^+^ cells in organoids at 2 or 4 weeks post-exposure to AD brain extracts or vehicle. Scale bars, 20 or 10 μm as indicated. (B) Representatives of AT8 and Thioflavin-S labeled hyperphosphorylated tau aggregates in organoids at 4 weeks post-exposure to AD brain extracts. Scale bars, 20, 10, or 5 μm as indicated. (C) Quantification of AT8 positive cells over time. n=4 independent experiments from 3 hPSC lines, each experiment used one hPSC line and contained 4-6 organoids. Data are presented as mean ± SEM. Unpaired t test with Welch’s correction, *p < 0.05, ns represents no significance. (D) Representatives of Gallyas silver staining of organoid at 4 weeks post-exposure to AD brain extracts or vehicle. Scale bar, 20 μm.

Confirmation of tau aggregate was further achieved through Thioflavin-S and AT8 co-staining, which marked Thioflavin-S+ and AT-8+ double-positive cells exclusively in the AD group (**Figure 3B**). Quantification revealed a substantial increase in AT8-positive areas in the AD group compared to the vehicle group (AD vs Vehicle, 0.88 ± 0.15 vs 0.10 ± 0.04) (**Figure 3C**), suggesting that AD brain extract was sufficient to induce tau pathology in organoids. To further confirm the formation of tau tangle-like structures, we performed Gallyas Silver Staining, a commonly used method for detecting tau tangles[57], and observed the formation of NFT-like structures in the AD group at four weeks post-exposure (**Figure 3D**). In all, our findings demonstrated that sAD brain extract can successfully induce tau pathology, including tau tangle-like aggregates, in healthy brain organoids.

### The AD neuroimmune organoids recapitulate neuroinflammation, phagocytosis of Aβ, and excessive microglial synaptic pruning

In the AD environment, accumulating amyloid aggregates trigger the activation of microglia and astrocytes, and stimulate neuroinflammation by releasing inflammatory factors[58–60]. Since neuroimmune organoids contain microglia and astrocytes, to determine whether neuroimmune organoids can replicate neuroinflammation in AD brains, we measured the mRNA expression of the pro-inflammatory cytokine, IL-6, and chemokine, CCL2, by qRT-PCR in the organoids at four to six weeks post the exposure of AD brain extracts or vehicle treatments. As shown in **Figure 4A**, compared to the vehicle group, the AD group exhibits a significant increase in mRNA levels of IL-6 (AD vs. Vehicle, 5.68 ± 0.37 vs. 1.03 ± 0.03) and CCL2 (AD vs. Vehicle, 1.46 ± 0.10 vs. 1.01 ± 0.01), demonstrating that AD neuroimmune organoids recapitulated neuroinflammation. In addition, microglia play a beneficial role in AD by phagocytosis and removal Aβ. To examine whether microglia in the AD group display this function, we conducted double staining of organoids with 6E10 and IBA1. 3D-reconstructive images from the Imaris software confirmed microglial engulfment of Aβ in the AD group, aligning with previous evidence of microglial involvement in Aβ clearance in AD[61, 62] (**Figure 4B**).

**Figure 4.**
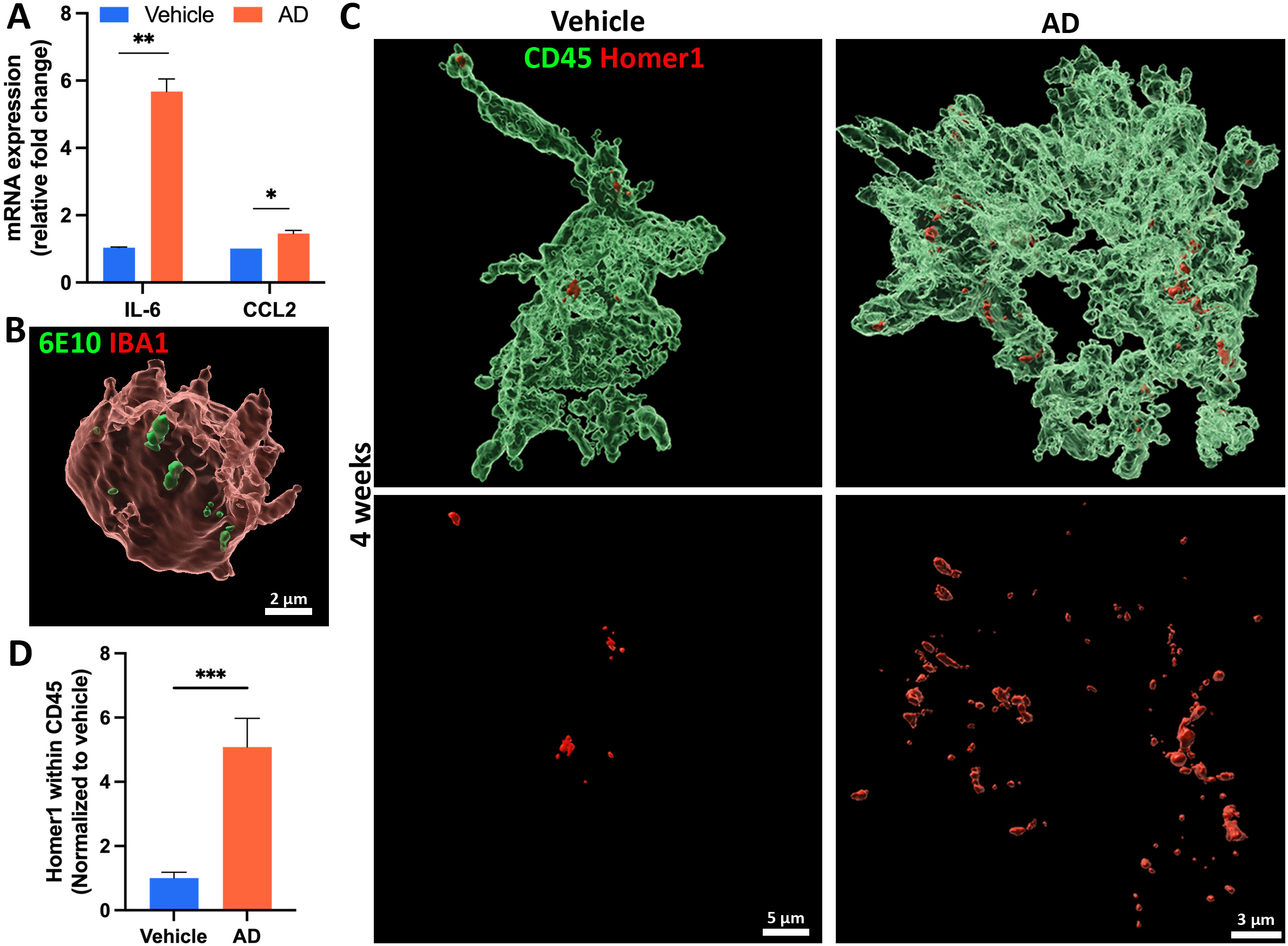
The AD neuroimmune organoids recapitulate neuroinflammation, phagocytosis of Aβ, and excessive microglial synaptic pruning. (A) qPCR analysis of mRNA expression level of *IL-6* and *CCL2* in organoids from 4 to 6 weeks post-exposure to AD brain extract or vehicle. n = 3 independent from two hPSCs lines, each experiment used one hPSC line and contained 4-6 organoids. Unpaired student’s t test, *p < 0.05, **p < 0.01. (B) 3D reconstructive image showing microglia phagocytizing Aβ. Organoid sections from the AD group treated with AD brain extracts were stained with IBA1 and 6E10. Scale bar, 2 μm. (C) 3D reconstructive image showing microglial synaptic pruning. Sections from organoids at 4 weeks post-exposure to the vehicle or AD brain extracts were stained with Homer1 and hCD45. Scale bars, 5 or 3 μm as indicated. (D) Quantification of Homer1 puncta engulfment within CD45-positive microglia for the vehicle and AD groups. The phagocytic activity of microglia was quantified by dividing Homer1 puncta volume with microglia volume. n = 4 independent experiments from three hPSC lines, each experiment used one hPSC line and contained 4-6 organoids. Data are presented as mean ± SEM. Unpaired t test with Welch’s correction, ***p < 0.001. Scale bars, as indicated.

One significant detrimental role of microglia in AD is the abnormal removal of synaptic materials[63]. We subsequently evaluated whether human microglia in AD neuroimmune organoids could recapitulate this phenomenon. Organoids were stained with CD45 and Homer1 (a post-synaptic marker) at four weeks post-exposure to AD brain extracts or vehicles. The 3D-reconstructed images revealed that microglia pruned synaptic materials (**Figure 4C**). Quantification of the volume of Homer1 puncta within the volume of CD45 indicated a significant increase of microglial pruning in the AD group (AD vs. Vehicle., 5.08 ± 0.9 vs. 1.00 ± 0.18) (**Figure 4D**). In summary, our results indicated that neuroimmune organoids can recapitulate neuroinflammation, as well as dynamic microglial functions in an AD-like 3D environment, which mirrors *in vivo* human microglia behaviors in the presence of AD pathologies.

### The AD neuroimmune organoids recapitulate synapse/ neuronal loss and impaired neural activity

Synapse/neuronal loss is the key feature of AD[6, 64] and AD animal models are often limited to recapitulating neuronal loss[65–67]. To investigate whether brain organoids challenged by AD brain extracts could recapitulate this process, we first examined synaptic integrity by staining organoids with Homer1. Quantitative analysis of the number of Homer1 puncta per 2500 μm^2^ revealed a reduction in the AD group (AD vs. Vehicle, 1336.30 ± 92.93 vs. 6013.48 ± 449.32) (**Figures 5A and 5B**), indicating synapse loss in the AD group. Subsequently, we stained organoids with active Caspase3 to assess neuronal death in organoids. Compared to the vehicle group, a significant increase in active Caspase3-expressing cells was detected in the AD group, demonstrating elevated neuronal death in the AD group (AD vs Vehicle, 14.11 ± 2.18 vs 2.66 ± 0.17) (**Figures 5C and 5D**).

**Figure 5.**
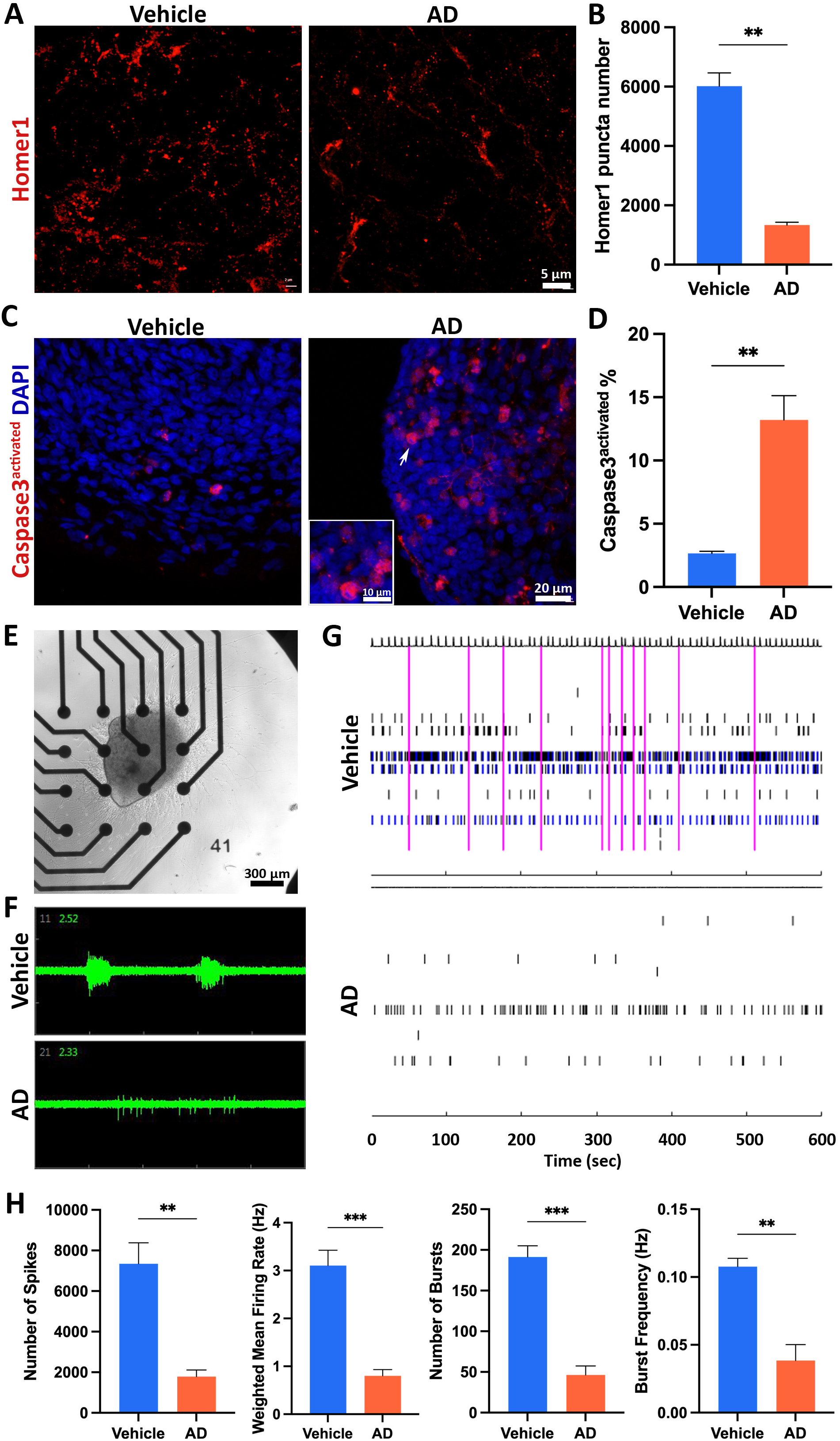
The AD neuroimmune organoids recapitulate synapse/ neuronal loss and impaired neural activity. (A) Representatives of Homer1 staining in the vehicle and AD groups at 4 weeks post-exposure to the vehicle or AD brain extracts, respectively. Scale bar, 5 μm. (B) Quantification of Homer1 puncta number per 2500 μm^2^. n = 4 independent experiments from three hPSC lines, each experiment used one hPSC line and contained 4-6 organoids. Data are presented as mean ± SEM. Unpaired t test with Welch’s correction, **p < 0.01. (C) Representatives of activated Caspase3 staining in the vehicle and AD groups at 4 weeks post-exposure to the vehicle or AD brain extracts, respectively. Scale bars, 20 or 10 μm as indicated. (D) Quantification of the percentage of Caspase3^+^ cells. n = 4, four independent experiments from three hPSC lines, each experiment used one hPSC line and contained 4-6 organoids. Data are presented as mean ± SEM. Unpaired t test with Welch’s correction, **p < 0.01. (E) Representative of organoid attached to MEA plate. Scale bar, 300 μm. (F) Representatives of spikes/bursts within 2 s on a single electrode in the organoids from the vehicle group or the AD group. (G) Representatives of spike raster plot generated from MEA recording raw data from the vehicle group and AD group. (H) Quantification of MEA parameters. n = 4 independent experiments from two hPSC lines, each experiment used one hPSC line and contained 4-6 organoids. Data are presented as mean ± SEM. Unpaired t test with Welch’s correction, **p < 0.01, ***p < 0.001.

To further understand whether synapse/neuronal loss causes functional defects, we assessed the neural activity in the organoids employing MEA assay (**Figures 5E to 5G**). Spontaneous neural activities were recorded and analyzed in organoids that were seeded on electrodes and cultured for two additional weeks after four weeks post-AD brain extract or vehicle treatment. Our data revealed a reduced number of spikes, mean firing rate, and number of bursts in the AD group (**Figure 5H**), suggestive of impaired neural activity in the AD group. Together, these results indicated an environment conducive to synapse/neuronal loss, accompanied by compromised neural activity in AD neuroimmune organoids.

### Anti-Aβ antibody Lecanemab relieved amyloid burden in AD neuroimmune organoids

Since AD organoids can recapitulate AD pathologies, we wonder whether the organoid model could be used for AD drug discovery. Lecanemab, an FDA-approved humanized monoclonal Aβ antibody for treating early AD, targets and neutralizes toxic Aβ, facilitating their clearance from the brain[68–70]. Given that our organoids displayed Aβ pathology after four weeks of AD brain extract exposure, we treated these neuroimmune organoids with Lecanemab for 2 consecutive weeks to evaluate their efficacy in our organoids. Staining with 6E10 and 4G8 post-treatment indicated that Lecanemab alleviated the amyloid burden in AD organoids (**Figures 6A and 6B**). It has been suggested that Lecanemab may induce phagocytosis and removal of Aβ by microglia in human brains[71, 72]. Therefore, we investigated the involvement of microglia in Lecanemab-mediated Aβ clearance in our organoids. Through staining with 6E10 and IBA1 and subsequent observation via 3D reconstructive images, we noted that microglia appeared to phagocytize Aβ in the organoids. Interestingly, Lecanemab-treated organoids exhibited increased phagocytosis of Aβ (**Figures 6C and 6D**). Overall, our results suggested that Lecanemab could relieve the Aβ burden in AD organoids, which might involve phagocytosis of Aβ by microglia. Consequently, our organoids demonstrated potential as a promising *in vitro* platform for testing AD therapeutics.

**Figure 6.**
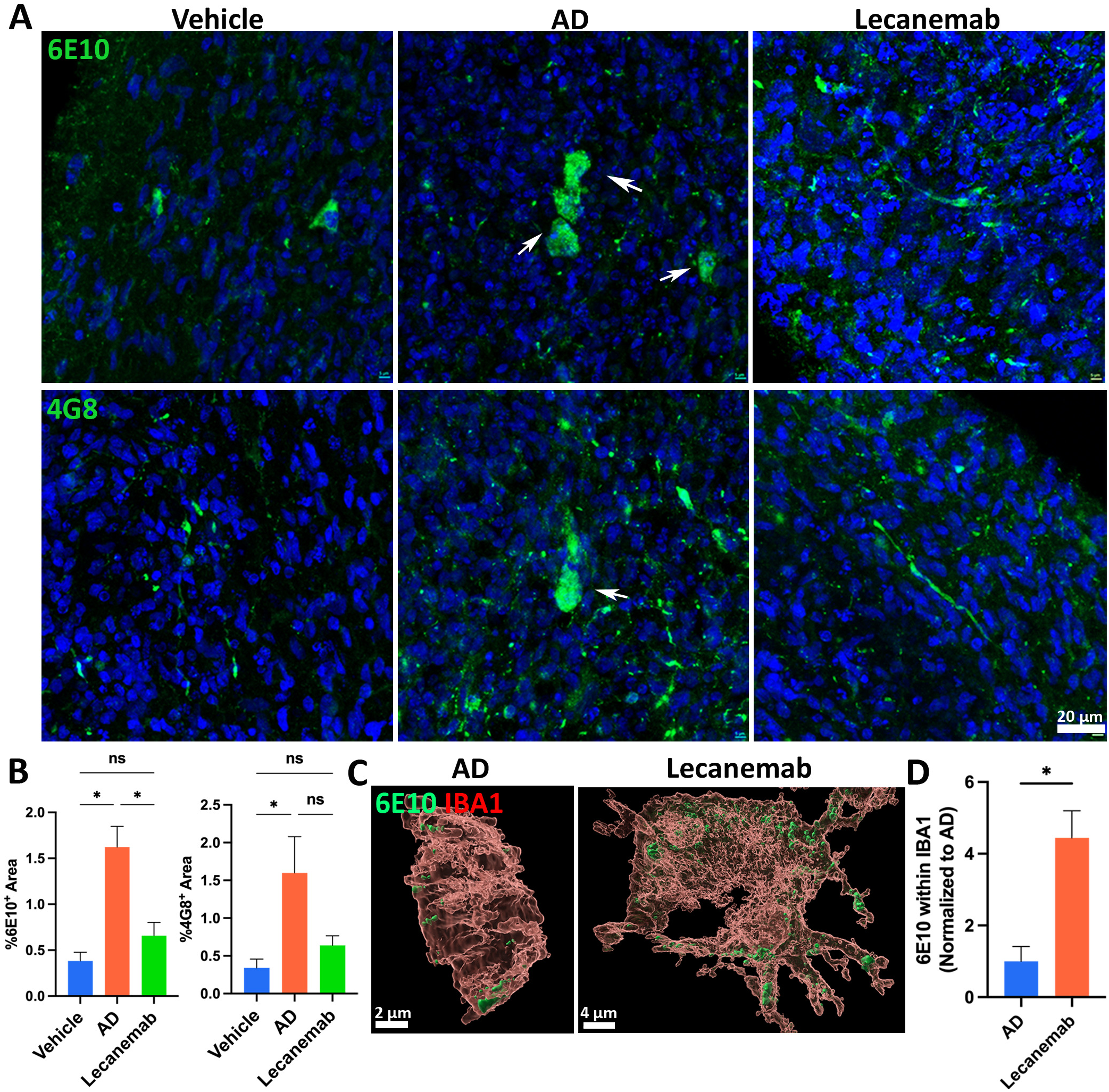
Anti-Aβ antibody Lecanemab relieved amyloid burden in AD neuroimmune organoids. (A) Representatives of 6E10 and 4G8 staining from the vehicle group, AD group, and Lecanemab-treated group. Scale bar, 20 μm. (B) Quantification of 6E10 and 4G8 positive area. n = 3, three independent experiments from three hPSC lines, each experiment used one hPSC line and contained 4-6 organoids. Data are presented as mean ± SEM. One-way ANOVA, *p < 0.05, ns represents no significance. (C) Representatives of 3D reconstructive figures showing the internalization of Aβ by microglia. Organoids from the AD group and Lecanemab-treated group were stained with IBA1 and 6E10. Scale bars, 4 or 2 μm as indicated. (D) Quantification of microglial phagocytosis of Aβ. n = 3 independent experiments from three hPSC lines, each experiment used one hPSC line and contained 4-6 organoids. Data are presented as mean ± SEM. Unpaired two-tailed t test with Welch’s correction, *p < 0.05.

## Discussion

Effectively modeling sAD to recapitulate multiple pathologies and developing platforms for drug discovery, have been challenged due to the complexity of the disease and the significant species differences between human and animal models. In this study, we presented a novel vascularized neuroimmune organoid model composed of human neurons, astrocytes, microglia, and blood vessels. Upon exposure to sAD individual-derived extracts, the organoid successfully recapitulated multiple pathological features in sAD, including Aβ plaque-like aggregates, neurofibrillary tangle-like aggregates, neuroinflammation, microglial synaptic pruning, synapse/neuronal loss, and impaired neural network. Furthermore, by validating the FDA-approved anti-Aβ antibody Lecanemab, the organoid model demonstrates its potential as a platform for *in vitro* AD drug development, particularly for immunotherapies.

In contrast to other hiPSC-based AD brain organoid models, our vascularized neuroimmune organoid model has significant advantages to enable us to more efficiently model AD and test AD therapeutics. (**1**) The new organoid model incorporates multiple key cell types that are affected in human AD brains under a pathophysiological relevant 3D human cell-centric environment. Our organoid simultaneously includes neurons, astrocytes, microglia, and vasculatures, which enables a more comprehensive understanding of cell-cell interactions and cell-type-associated pathological events in AD, such as neuroinflammation and microglial synaptic pruning, which are absent in most AD organoid models (**Figure 4**). (**2**) The new organoid model can effectively develop multiple AD pathologies within a relatively short time frame, suitable for studying AD, particularly sAD. Previous AD organoid models were usually derived from fAD iPSC lines, which need long-term culture to replicate the AD pathologies. For instance, previous studies indicated that it took at least 5 months to exhibit amyloid plaque-like and neurofibrillary tangles-like aggregates[34], or 90 days to observe Aβ aggregates and hyperphosphorylated tau protein[29], in fAD iPSC-derived brain organoids. Additionally, a recent study using healthy iPSC lines-derived organoids to model sAD through serum exposure required 3 to 4 months of culture to recapitulate Aβ aggregates and hyperphosphorylated tau [36]. However, in our organoids, 4 weeks post-exposure to AD brain extracts, we successfully recapitulated multiple relatively mature AD pathologies, including Aβ plaques-like aggregates and tau tangles-like aggregates, neuroinflammation, synapse/neuronal loss, in around 1.5-month-old organoids. This suggests that AD brain extracts can serve as a strong trigger to induce AD pathologies for studying AD, particularly sAD. In addition, complex cell types, particularly microglia and related neuroinflammation, absent in the previously mentioned organoids, may also contribute to efficiently recapitulating the AD pathologies in our organoids. With a similar strategy, it is possible to efficiently model fAD by exposing fAD-derived brain extracts to fAD iPSC-derived organoids. (**3**) The new organoid model is useful to serve as an *in vitro* platform to advance AD drug development, particularly antibody-based therapeutics (**Figure 6**), because the organoids contain microglia and can efficiently recapitulate multiple AD pathologies, and microglia are expected to be a major cell component to degrade the antibody-binding protein aggregates, such as Aβ.

Our results demonstrate that AD brain extracts can efficiently induce multiple AD pathologies in organoids, particularly Aβ and tau pathologies. To induce AD pathologies, we utilized brain extracts from sAD individual postmortem brain tissues and hypothesized that the Aβ and tau seeds in the brain extracts could induce multiple pathologies. As expected, post-exposure to sAD brain extracts, brain organoids can efficiently recapitulate Aβ and tau pathologies. We first observed intraneuronal Aβ accumulation, as indicated by punctate-like high-intensity 6E10-positive signals (**Figure 2B**), at 2 weeks postAD brain extract exposure. By the fourth week, these accumulations had progressed to extracellular Aβ plaque-like structures, indicated by the colocalization of 6E10 and Thioflavin-S (**Figures 2B, 2C, and 2F**), suggesting a developing pattern from intraneuronal aggregation to extracellular deposition. This pattern is in line with the findings in AD and Down Syndrome individuals, where intraneuronal amyloid aggregates were found in brain regions vulnerable to dementia, such as the hippocampus and entorhinal cortex, preceding the formation of extracellular plaques[73–77]. Similarly, organoids demonstrated puncta-like aggregates of hyperphosphorylated tau 2 weeks post AD brain extract exposure and exhibited tangle-like structures by the fourth week post-exposure, indicated by the Thioflavin-S and Gallyas Silver Staining (**Figures 3 A, 3B, and 3D**).

On the other hand, our results provide direct evidence that proteopathic seeds, not only Aβ but tau, can spread among human cells and tissue-like 3D structures, and induce AD-like Aβ and tau pathologies. This aligns with the clinical observations that Aβ contaminated c-hGH can cause Aβ pathologies and CAA in humans over decades[49, 50, 58]. This also corroborates the concept of the prion-like property of AD proteopathic proteins and the possible transmission of AD between humans by accidents in medical or surgical procedures, raising the awareness to prevent iatrogenic human transmission of AD.

Neuroinflammation and synapse/neuronal loss are known to appear along the progression of AD[78]. In the early stages of the disease, stimuli like extracellular Aβ activate neuroinflammatory responses that may recruit microglia for Aβ clearance[58, 59]. Consistent with this, in our organoids, mRNA expression levels of inflammatory cytokines in the AD group (**Figure 4A**). Furthermore, Aβ signals were detected inside of microglia, indicating microglial phagocytosis and clearance of Aβ in organoids (**Figure 4B**). Additionally, align with previous studies showing that excessive microglia-mediated synaptic pruning in AD animal models [63, 79], we observed a significant microglia synaptic removal (**Figures 4C, 4D**), and a significant synaptic materials decrease in AD organoids (**Figures 5 A and 5B**). Additionally, elevated neuronal death was confirmed by activated Caspase3 signaling increment in the AD group (**Figures 5 C and 5D**). Although studies have found neuronal hyperactivity in the presence of amyloid beta pathology, *in vivo* research has proved that tau dominated Aβ when both were present in transgenic mice models[80]. In agreement with these observations, our MEA assay of organoids revealed impaired neuronal networks (**Figures 5E to 5H**), with fewer spikes and bursts recorded compared to the vehicle group, corroborating previous research findings.

The neuroimmune organoids, containing multiple cell types (including microglia) and exhibiting AD-like pathologies, are particularly valuable for testing AD therapeutics, especially immunotherapies. The recent FDA-approved anti-Aβ monoclonal antibody, Lecanemab, underscores the tremendous potential of antibody-based immunotherapies for tackling AD[5, 81, 82]. However, species differences may hinder the translational efficacy of preclinical animal models in clinical trials, highlighting the urgent need for human models to test these immunotherapies before clinical trials. (**1**) Recent cryo-EM findings have uncovered species-specific aggregate structures of proteopathic proteins in AD[83]. Aggregate structures observed in AD animal models are often not present in individuals with AD[84]. This discrepancy potentially impedes our assessment of the efficacy of the therapeutics and contributes to translation failures. (**2**) Antibodies-based therapeutics targeting toxic proteins, such as targeting Aβ, are usually tested in animals. Before moving to clinical trials, successful antibody candidates undergo a humanization process to avoid immune rejection, such as by replacing the murine IgG with the human version [85]. However, the humanization process may substantially compromise the efficacy of the antibodies and even cause side effects[86]. Thus, direct testing of these human antibodies in human models is crucial. (**3**) Antibody-targeted protein aggregates, such as Lecanemab-binding Aβ protofibrils, are hypothesized to be recognized and removed by microglia, the immune cells in the brain[70–72]. However, there are no suitable human models available that can recapitulate AD pathologies and contain human microglia for directly testing these human versions of antibody-based therapeutics. Our organoids contain multiple cell types, including microglia, and develop AD-like pathologies that offer a unique platform to solve this limitation. To assess whether our organoid model could serve as an ideal *in vitro* platform for testing AD drugs, we validated Lecanemab in our organoids. Following a 4-week post-exposure to AD brain extracts, we treated our AD organoids with Lecanemab for two weeks and observed a significant decrease in amyloid burden. Notably, we also observed an increased microglial engulfment of Aβ in the Lecanemab-treated group (**Figure 6**), providing direct evidence that human microglia involved the Lecanemab-mediated Aβ removal.

While the inclusion of blood vessels improves the overall health of the organoids to better model the complexity of the brain environment, we did not detect the presence of CAA, where Aβ fibrils are deposited along the cerebral vasculatures, a common vascular pathology in AD[64]. This may require an extended culture period post-AD extract exposure. In addition, a critical challenge for iPSC-based brain organoid models is the relatively immature state of human cells[87]. Recent studies, including ours, have developed chimeric brain models by transplanting iPSC-derived organoids or neural cells into animal brains, where human cells can survive and mature as animals age under the conducive physiological *in vivo* environment[44, 45, 88, 89]. It will be promising to transplant our organoids into animal brains and test whether our organoids can better model AD features in the context of aged human cells within the host brain in the future. Overall, our innovative vascularized neuroimmune organoids present unique opportunities to model sAD and advance drug discovery for AD.

## Material and Methods

### Culture and quality control of hPSC lines

Three different hiPSC lines, KOLF2.1, ND2, and GCaMP, as well as one hESC cell line CAGG, were used in this study (**Supplementary Table S1**). The hiPSCs were maintained under feeder-free condition and cultured on hESC-qualified Matrigel (Corning) coated dish in mTeSR plus media (STEMCELL Technologies). The hiPSCs were passaged at approximately 70% confluency with ReLeSR media (STEMCELL Technologies).

### pNPC generation and culture

Small molecule-based protocols were applied when generating pNPCs as in our previous studies[45, 90]. Neural differentiation in the embryoid bodies (EBs) was induced by dual inhibition of SMAD signaling. In summary, EBs were cultured in neural induction medium composed of DMEM/F12 (HyClone) and 1 × N2 Supplement (Thermo Fisher Scientific) supplied with inhibitor SB431542 (2 μM, Stemgent) and noggin (50 ng/mL, Peprotech) for 6 days before plating on growth factor-reduced Matrigel (BD Biosciences) coated plates. EBs were cultured with neural induction medium supplied with laminin (1 μg/mL, Corning) for 7 days, and then pNPCs in the form of neural rosette were manually isolated from surrounding cells. Isolated pNPCs were expanded for 5 to 7 days, depending on the confluency, in pNPC medium, which is composed of a 1:1 mixture of Neurobasal (Thermo Fisher Scientific) and DMEM/F12, supplemented with 1×N2, 1×B27-RA (Thermo Fisher Scientific), FGF2 (20 ng/mL, Peprotech), human leukemia inhibitory factor (hLIF, 10 ng/mL, Millipore), CHIR99021 (3 μM, Biogems), SB431542 (2 μM), and ROCK inhibitor Y-27632 (10 μM, Tocris).

### PMP generation and culture

PMPs were generated as previous studies[44, 91]. Briefly, embryoid bodies (EBs) were generated from hiPSCs and induced to yolk sac EBs (YS-EBs) by adding bone morphogenetic protein 4 (BMP4, 50 ng/mL, Peprotech; to induce mesoderm), vascular endothelial growth factor (VEGF, 50 ng/mL, Peprotech; endothelial precursors), and stem cell factor (SCF, 20 ng/mL, Miltenyi Biotech; Hematopoietic precursors) into mTeSR1 media (STEMCELL Technologies) for 5 days. Next, YS-EBs were plated into dishes and cultured with PMP medium composed of interleukin-3 (IL-3, 25 ng/mL, Peprotech) and macrophage colony-stimulating factor (M-CSF, 100ng/mL, Peprotech) in XVIVO media (Lonza). Human PMPs emerged into the supernatant 2 to 3 weeks after plating and were continuously produced for more than 3 months.

### VP generation and culture

VPs were generated using a published protocol with minor modifications[46]. In brief, hiPSCs were dissociated to form aggregates in a low-attachment plate and differentiated into mesoderm with CHIR99021 (12 μM, Peprotech) and BMP4 (30 ng/mL) in a 1:1 mixture of Neurobasal and DMEM/F12, supplemented with 1×N2, 1×B27-RA for 3 days. Then, vascular lineage induction was performed with VEGF (100 ng/mL) and forskolin (2 μM) for 2 days. Cell aggregates were then embedded into growth factor-reduced Matrigel, cultured with VEGF (100 mg/mL), FGF2 (100 ng/mL), and 15% FBS (Gibco) in XVIVO media for 5 days to allow vessel sprouting. Cell aggregates were dissociated by TrypLE into single cells, which served as the VP for the following organoid generation.

### Brain extract preparation and tau seeding activity confirmation

To obtain 10% (w/v) brain extracts, frozen temporal cortex tissues of histopathologically confirmed AD cases with Braak Stages V-VI from the Brain Tissue Resource Center, McLean Hospital, Belmont, MA, USA, were prepared in homogenization buffer (20 mM Tris–HCl, pH 8.0, 0.32 M sucrose, 10 mMβ-mercaptoethanol (β-ME), 5 mM MgSO 4, 1 mM EDTA,10 mM glycerophosphate, 1 mMNa 3VO4, 50 mM NaF,2 mM benzamidine, 1 mM 4-(2-aminoethyl) benzenesul-fonyl fluoride hydrochloride (AEBSF), and 10 μg/ml each aprotinin, leupeptin, and pepstatin), and centrifuged at 10,000×g for 30 min, as previously reported[92, 93]. Tau seeding activity of brain extracts was confirmed (**Figure S5**), using *in vitro* seeded tau aggregation assay, in which HEK-293T cells expressing HA-tau_151-391_ was treated with brain extracts and lysed in RIPA buffer. Seeded HA-tau_151-391_ aggregates were yielded by ultracentrifugation and analyzed by immuno-blots, as previously reported[92, 93].

### Organoid assembly, culture, brain extract treatment, and drug treatment

To generate vascularized immune-brain organoids, 30,000 pNPCs, 12,000 PMPs, and 7,000 VPs were co-cultured for organoid self-assembly. Organoids were cultured in a 1:1 mixture of NPC medium (1:1 mixture of Neurobasal and DMEM/F12, supplemented with 1×N2, 1×B27-RA, FGF2 (20 ng/mL)) and PMP medium for 5 days to allow proliferation, and were transferred to ND medium, which is composed of a 1:1 mixture of Neurobasal and DMEM/F12, supplemented with 1×N2, 1×B27-RA, BDNF (10 ng/mL, Peprotech), GDNF (10 ng/mL, Peprotech), L-Ascrobic acid (200 nM, Sigma Aldrich), c-AMP (1 μM, Sigma Aldrich), IL-34 (100 ng/mL, Peprotech), M-CSF (25 ng/mL), TGF-β1 (50 ng/mL, Peprotech). After 10 days of differentiation, vehicle or brain extracts derived from AD patients, or age-matched healthy individuals (10 μL/mL), were added to medium for two days. The organoids were collected at 2 weeks or 4 weeks post-exposure to brain extracts or the vehicle. Four weeks post exposure to brain extracts, Lecanemab (Sellekchem, Cat#A3112) was added to medium at a concentration of 10 mg/mL. Samples were collected after 2 weeks of Lecanemab treatment.

### Immunohistochemistry and cell counting

Organoids were fixed with 4% paraformaldehyde and then cryo-sectioned at 15 μm thickness for immunostaining. The tissues were blocked with a blocking solution (5% goat or donkey serum in PBS with 0.2% Triton X-100) at room temperature for 1 hr. Primary antibodies were diluted in the same blocking solution and incubated at 4 °C overnight. Sections were washed with PBS and incubated with secondary antibodies for 1 hr at room temperature. Sections were then washed with PBS and were mounted with anti-fade Fluoromount-G medium containing 1,40,6-diamidino-2-phenylindole dihydrochloride (DAPI) (Southern Biotechnology). Antibodies used were listed in **Supplementary Table S2**. For Thioflavin-S staining, sections were washed with 70% and 80% ethanol for 1 minute, respectively, followed by 15 minutes of incubation of 0.05% freshly prepared and filtered Thioflavin-S solution. Samples were then washed with 80% and 70% ethanol sequentially for 1 minute, and finally rinsed with water. Images were captured with a Zeiss LSM 900 confocal microscopy. Image analysis was performed using Fiji (NIH). Relative fluorescence intensity was presented as normalized value to the vehicle group. Cells, plaques, or synapses were counted with Fiji. At least three fields of each organoid were chosen randomly to count after Z projection. The data are repeated three to four times (n=3 or 4) from 3 hPSC lines, each experiment used one hPSC line and contained 4-6 organoids.

### Gallyas Silver Staining

The Gallyas silver staining method was used to stain the Tau tangles (57), details are included in Supplementary materials. Slides were washed in distilled water for one minute and transferred immediately to alkali silver iodide for 15 minutes. Next, slides were washed for 1 minute in 2% oxalic acid, followed by three washes in distilled water for 1 minute. Then the slides were incubated in alkaline silver iodide solution for 2-5 minutes followed by three rinsing in 0.5% acetic acid for 1 minute each. Then slides were incubated in physical developer solution for 15 -20 minutes. The developer was made fresh before use in a 1:1 ratio (solution A: solution B). Next, the samples were washed in 0.5% acetic acid and 1% sodium thiosulfate for 5 minutes each. Subsequently, samples were washed in distilled water for five minutes, and incubated in 0.5% gold chloride solution, and sodium thiosulfate for 5 minutes each. Finally, the samples were washed in distilled water for five minutes, counterstained with nuclear fast red, and mounted using Permount solution.

### RNA isolation and qPCR

Total RNA isolation was performed with TRIzol Reagent (Invitrogen) and complementary DNA was prepared with SuperScript IV First-Strand Synthesis System (Invitrogen)[94]. The qPCR assays were performed with SYBR Green PCR Master Mix in QuantStudio 3 (Applied Biosystems), primers used were listed in **Supplementary Table S3**[95]. The 2^−ΔΔCt^ method was used to calculate relative gene expression after normalization to the β-actin internal control.

### Microelectrode Arrays (MEA)

Organoids at around 21 days were seeded into 48-well transparent MEA plates (Axion Biosystems) at one organoid per well. MEA assays were performed with Maestro Pro platform (Axion Biosystems) and recorded using AxIS software. Organoids were fed with BrainPhys media (STEMCELL Technologies) supplemented with 1×N2, 1×B27, 20 ng/mL BDNF, 20 ng/mL GDNF, as well as brain extract or vehicle. For recording, following a 5 min resting time in the instrument, each plate was recorded for 10 min to calculate the spike per well. Experiments were repeated for four times, each experiment used one hPSC line and contained 3-4 organoids. MEA analysis was performed using the Axion Biosystems NeuralMetrics Tool.

### Statistics and reproducibility

All data represent mean ± SEM. Significance is determined using a two-tailed unpaired t-test with Welch’s correction for comparing two independent groups, or a one-way ANOVA test with Bonferroni post-hoc test for comparing three or more groups. A p value < 0.05 was considered significant. Analyses were performed with GraphPad Prism 10. All experiments were independently performed at least three times.

## Acknowledgments

This work was in part supported by grants from the lab startup package, the Showalter grant, and the Purdue Institute for Integrative Neuroscience research grant (iPSC/Organoid Technologies) to X.R. We appreciate Ying Liu from Florida International University for providing ND2 iPSC and CAGG hESC lines in this study. We thank Xinqi Guo and Raymond Pan from the Xu Laboratory for their assistance with immunohistochemistry.

## Authors’ contributions

R.X. and J. Y designed the experiments and interpreted the data. J.Y. carried out most of the experiments with technical assistance from C. X., C.E., J.C.M, and L.F, prepared the brain extracts with quality control. C. X., and Y. Y assistance with MEA recording. L.F., R. JC., Y. C., and Y. Y. provided critical suggestions. R.X. directed the project and wrote the manuscript together with J.Y. with input from all co-authors.

